# Epigenetic age acceleration predicts subject-specific white matter degeneration in the human brain

**DOI:** 10.1101/2022.11.14.516491

**Authors:** Benjamin T. Newman, Joshua S. Danoff, Morgan E. Lynch, Stephanie N. Giamberardino, Simon G. Gregory, Jessica J. Connelly, T. Jason Druzgal, James P. Morris

**Author notes:** These authors are co-senior authors of this work (T. Jason Druzgal, James P. Morris).

## Abstract

Epigenetic clocks provide powerful tools for estimating health and lifespan but their ability to predict brain degeneration and neuronal damage during the aging process is unknown. In this study, we use GrimAge, an epigenetic clock correlated to several blood plasma proteins, to longitudinally investigate brain cellular microstructure in axonal white matter from a cohort of healthy aging individuals. Given the blood plasma correlations used to develop GrimAge, a specific focus was made on white matter hyperintensities, a visible neurological manifestation of small vessel disease, and the axonal pathways throughout each individual’s brain affected by their unique white matter hyperintensity location and volume. 98 subjects over 55 years of age were scanned at baseline with 41 returning for a follow-up scan 2 years later. Using diffusion MRI lesionometry, we reconstructed subject-specific networks of affected axonal tracts and examined the diffusion cellular microstructure composition of these areas, both at baseline and longitudinally, for evidence of cellular degeneration. A chronological age-adjusted version of GrimAge was significantly correlated with baseline WMH volume and markers of neuronal decline, indicated by increased extracellular free water, increased intracellular signal, and decreased axonal signal within WMH. By isolating subject-specific axonal regions ‘lesioned’ by crossing through a WMH, age-adjusted GrimAge was also able to predict longitudinal development of similar patterns of neuronal decline throughout the brain. This study is the first to establish a relationship between accelerated epigenetic GrimAge and brain cellular microstructure in humans.

## Introduction

Recent advances in epigenetic sequencing and analysis have led to the development of a number of epigenetic clocks that act as biomarkers for chronological age^1,2^. GrimAge is an epigenetic clock that calculates expected time-to-death due to all-cause mortality based on a number of surrogate DNA methylation based biomarkers of stress and physiological risk^3^.

AgeAccelGrim is an age-adjusted version of GrimAge shown to be highly predictive of time-to-coronary heart disease, congestive heart failure, hypertension, type 2 diabetes, and physical functioning^3^. It is well established that cardiovascular health affects brain integrity and cognitive functioning^4–6^ and has been implicated in the formation of white matter hyperintensities (WMH) on T2-weighted brain MRI. WHM are thought to be the consequence of small vessel disease (SVD) which can cause microinfarcts, edema, and cortical thinning^7–9^. WMH are associated with declining cognitive and perceptual functioning and are thus an important marker for age-related brain health^10,11^. In this study, we pair AgeAccelGrim with an advanced diffusion microstructure analysis technique, 3-Tissue Constrained Spherical Deconvolution (3T-CSD), in a subject-specific manner to test associations between AgeAccelGrim estimates of mortality risk and brain cellular microstructure. Analyses were focused on WMH due to their connection to SVD, as well as on the ability of AgeAccelGrim to predict future neuronal decline in an aging cohort.

At the population level, cardiovascular risk factors in humans have been strongly linked to degenerative changes in brain structure using brain diffusion MRI (dMRI). Greater arterial stiffness has been associated with reduced white matter (WM) fractional anisotropy in the corpus callosum and corona radiata as well as lower grey matter (GM) density in the thalamus^12^. In a recent study using dMRI and positron emission tomography (PET), MRI-based markers of SVD (measured by WMH volume) were more correlated with dMRI extracellular free water than PET measures of Tau or Aβ, and SVD contributed more to diffusion alterations than did biomarkers of Alzheimer’s disease^7^. Another dMRI study suggested that diffusion differences between patients with SVD and healthy controls were primarily driven by increased extracellular free water rather than neuronal tissue alterations, and that this finding predicted clinical status^13^. These dMRI-based degenerative changes follow similar trajectories across the ‘healthy’ population (those explicitly without a diagnosed neurodegenerative disease), but there is wide individual variation in the chronological age at which degenerative brain changes begin and how they progress.

The GrimAge model was generated from the Framingham Heart Study Offspring Cohort^14^ and is specifically composed of 12 DNA methylation (DNAm) based biomarkers for plasma proteins, plus age, gender, and smoking pack-years, regressed to time-to-death. The age-adjusted version of GrimAge presents a powerful means for studying the effects of cardiovascular health on the brain. Not only is the clock optimized to estimate plasma proteins sensitive to aging and general heart health, but by establishing DNAm correlates for factors such as smoking pack-years, GrimAge allows for the assessment of levels of methylation present on the relevant site even in the absence of self-report metrics or any history of smoking at all. Age-adjusted GrimAge is not only reported to be more closely correlated with actual time-to-death than self-reported smoking pack-years alone, but it allows for risk evaluation and stratification of all non-smokers, providing a more powerful and accurate statistical sample^3^.

GrimAge may be a useful biomarker for neuronal decline related to WMH and SVD. WMH volume is one of the few neuroimaging markers of SVD that has been associated with AgeAccelGrim,^15^ though no conclusive link between AgeAccelGrim and WMH has been established^16–18^. The difficulty in establishing this link highlights the shortcomings of whole brain or one-size-fits-all approaches to studying features such as WMH that vary greatly in presentation between individuals. Individual differences in age-related neuronal decline are evident in whole-brain metrics and particularly in the presentation of WMH. Analyzing the composition of WMH presents a challenge for typical neuroimaging analysis pipelines but particularly using 3T-CSD. Each subject has a different spatial location of WMH volume and a distinct spatial progression longitudinally. The location of WMH has been implicated in cognitive deficits in patients with SVD^19^ so it is crucial to account for subject-specific WMH features. Being located in the predominantly axonal WM, WMH can induce damage and Wallerian degeneration on axonal pathways traversing the lesioned area^9^ presenting a challenge to investigate distant regions of the brain that may show signs of damage that form unique subject-specific spatial patterns.

To address this problem of subject-specific spatial patterns, our study applied 3T-CSD microstructural metrics within a lesionometry framework^20,21^. 3T-CSD analysis of cellular microstructure has previously been used to successfully characterize WMH and is also able to detect areas of ‘normal appearing’ WM that develop into WMH following cerebrovascular injury^22,23^. Lesionometry is a fusion of voxel-wise diffusion metrics and subject specific lesion analysis to isolates axonal networks that traverse lesioned voxels with diffusion tractography and analyze the cellular microstructure within that network (Figure 1). This technique has previously been applied to the study of multiple sclerosis, and was able to uncover a number of diffusion metrics and network measures, such as the proportion of lesioned volume to whole affected network volume referred to as ‘lesion load’, to cognitive, learning, and memory symptoms^21^.

**Figure 1:**
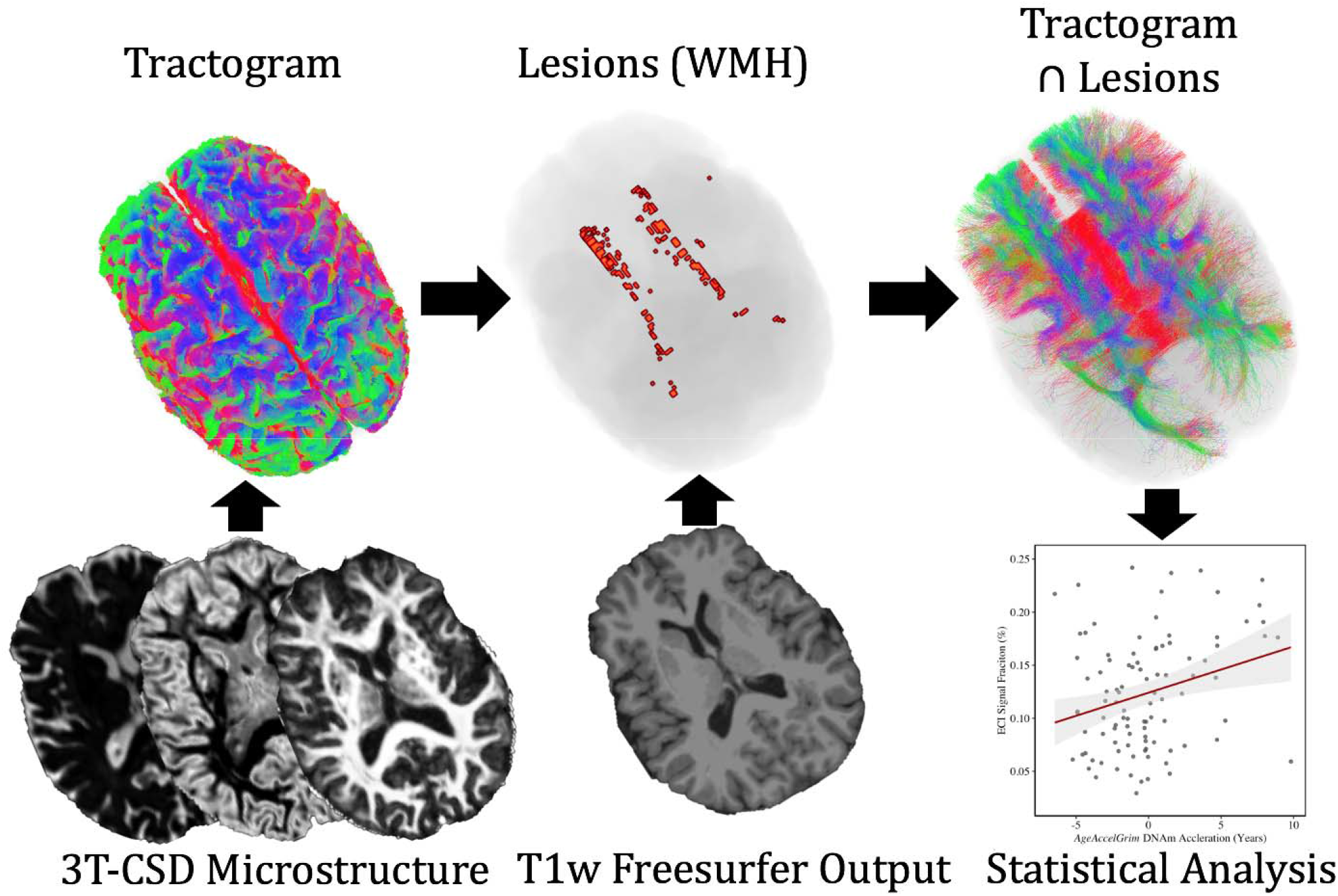
Flowchart demonstrating Lesionometry analysis pipeline, first described in Chamberland et al. (2020) and Winter et al. (2021). Subjects dMRI data is processed thorough the 3T-CSD microstructure pipeline and FODs are used to generate a whole brain tractogram. ‘Lesions’ representing WMH are derived from T1-weighted images taken from the ‘recon-all’ processing pipeline in freesurfer. Voxels traversed by tracts that also traverse WMH lesions are included for final microstructure statistical analysis.

Using this subject-specific method to focus highly sensitive 3T-CSD microstructural methods may be the key to finding a relationship between GrimAge clock measurements and changes in brain microstructure. Due to its connection to cardiovascular risk factors, GrimAge may be well-suited to determining where an individual falls on the age-related decline trajectory. This study aims to evaluate GrimAge as an effective peripheral blood indicator of SVD-driven damage to the brain.

## Methods

### Subjects

Participants were recruited from the ongoing Virginia Cognitive Aging Project (VCAP), a multi-year cross-sectional and longitudinal study of cognition in over 5,000 participants^24,25^. VCAP subjects have been recruited from the local community and have agreed to participate in multiple study visits over several years. A subset of older adult subjects (ages 58-81) was selected from VCAP for the present study, and an equal number of subjects from each quantile of performance based on cognitive tasks during prior visits was recruited for additional MR imaging. This age range was selected to target the prodromal phase of neurodegeneration, to test predictive biomarkers of cognitive decline. Additionally, we recruited equally across the sample by cognitive performance to have an adequately varied sample that is representative of the population.

98 subjects were recruited for baseline neuroimaging with an age range between 58-81 (mean = 68 ± 5.67 S.D.) years old. There were 30 male and 68 female participants with average ages of 67.53 ± 5.44 S.D. and 68.84 ± 5.75 S.D., respectively. Follow-up neuroimaging planned for 1 year was delayed by the COVID-19 pandemic and ended up occurring approximately 2 years later. Many of the elderly subjects declined follow-up neuroimaging due to ongoing COVID-related concerns. A total of 41 subjects were successfully recruited for follow-up scans with an age range of 61-81 (mean = 69 ± 5.00 S.D.) years old. This follow-up group was composed of 16 male and 25 female participants with average ages of 68.19 ± 4.96 S.D. and 69.86 ± 5.02 S.D., respectively.

### Image Acquisition

All subjects were scanned at the University of Virginia using a Siemens Prisma 3T MRI with a 32-channel head coil. T1-weighted images were acquired using the ADNI3 designed MP-RAGE sequence^26^ with an isotropic voxel size 1.0×1.0×1.0mm^3^, TE=2980ms and TR=2300ms with full dimensions of a 208×240×256 viewing window. Diffusion-weighted images were acquired with an isotropic voxel size of 1.7×1.7×1.7mm^3^, TE=70ms and TR=2900ms; using a multi-shell protocol, 10 b=0 images and 64 gradient directions were collected at both b=1500s/mm^2^ and b=3000s/mm^2^. An identical imaging protocol was used at both baseline and follow-up.

### Preprocessing

Each diffusion image set was analyzed using SS3T-CSD^27,28^ implemented in the open source software MRtrix and MRtrix3Tissue^27,29^. Several preprocessing steps utilized FSL^30,31^. Diffusion images were denoised^32^, corrected for Gibbs ringing^33^, susceptibility distortions^31^, subject motion^34^, and eddy currents^35^. Skull-stripping was performed and volumetric data was gathered by analyzing each subject’s T1 image from each visit using the ‘recon-all’ pipeline in Freesurfer version 6.0.1^36^. All images were upsampled to 1.3⨉1.3⨉1.3mm and a whole brain mask was derived by rigidly registering each subject’s skull-stripped T1 image from the appropriate scanning session to the average b=0 s/mm^2^ dMRI acquisition using ANTs^37^.

Response functions were generated^38^ from each tissue type (WM, GM, and CSF) and the white matter fiber orientation distribution (FOD) was then resolved at the voxel-wise level by processing the outermost b-value shell (b=3000s/mm^2^) using single-shell constrained spherical deconvolution, a technique to separate directional axonal signal from intracellular and extracellular isotropic diffusion^27^. Primarily for the purposes of visualization, a cohort-specific FOD template was generated from 20 subject’s WM FODs acquired from both baseline and follow-up for a total of 40 FOD images being used in template construction. Each subject was subsequently registered to this template using the individual WM FODs from each timepoint to warp tractography and signal fraction maps into a common space^39^. However, all calculations used in the analysis were derived from native acquisition space images.

### Tractography

Probabilistic tractography was performed on each subject’s diffusion images by applying the iFOD2 algorithm which propagates streamlines between voxels based on the direction and amplitude of the underlying WM FOD^40^. Seeding of streamlines was performed by randomly selecting voxels within the whole brain mask. Streamlines were seeded and generated until 10,000,000 tracts were created that were each longer than 2.6mm without terminating. These streamlines were then pruned to 2,000,000 total tracts using spherical-deconvolution informed filtering of tractograms (SIFT), which ties the number of streamlines in each voxel to the magnitude of the underlying FOD^41^. This process matches the randomly generated streamlines to the underlying anatomically derived signal and prevents a biologically implausible number of tracts from traversing the same voxel. As all subjects provide the same number of total streamlines, the number of tracts inferences can be made in a between-subject or longitudinal manner.

### Diffusion Microstructure

3T-CSD measurements of brain cellular microstructure were calculated directly from each subject’s FODs at each timepoint. 3T-CSD is a voxel-wise quantitative method that measures cellular microstructure within each voxel fitting into intracellular anisotropic (ICA, WM-like), intracellular isotropic (ICI, GM-like), and extracellular isotropic (ECI, CSF-like/Free Water) compartments^42^. This approach allows for contributions from each cellular microstructure compartment to be calculated for each image in any defined ROI. The quantitative measurement of 3 different tissue compartments, the improvements to tractography by separating isotropic signal from anisotropic WM for tractography, and the specific ability to measure extracellular freely diffusing water (a potentially highly sensitive marker for neuronal degeneration during aging^43^).

### Identification of WMH

Voxels composing WMH were identified through application of the ‘recon-all’ pipeline to each subject’s T1 images at each timepoint collected. Freesurfer identifies WMH via segmentation of WM voxels followed by a voxel-wise probabilistic local and intensity-related analysis informed by a library of manually segmented images^36^. While Freesurfer has been shown to systematically underestimate WMH volume when applied to T1 images, volume estimations were closely correlated with Fazekas score, a measure of WMH severity^44^. No FLAIR acquisition was collected during this study. Masks highlighting voxels containing WMH were rigidly transformed into the space of the respective diffusion image acquired during the same session using the previously generated ANTs rigid transform from the T1 to average b=0 registration described previously^37^.

### Lesionometry

Lesionometry was used to examine the relationship between WMH volume, cellular microstructure composition, and spatial positioning to whole brain structure. This recently developed technique was applied to generate subject- and timepoint-specific measures of ‘lesion-load’^20,21^. This method examines microstructural metrics within voxels traversed by WM fiber bundles, and theoretically the axons they model, that also traverse WMH (Fig. 1). WMH masks generated by Freesurfer and registered to the diffusion space were used to filter tracts from the final whole brain tractogram after the processing pipeline described earlier. To reduce spurious individual tracts from being overrepresented in analysis a threshold was applied so that voxels were only included in the final ROI if at least 10 separate tracts traversed both the voxel and a WMH. Volumetric and 3T-CSD measurements were then measured within each ROI from all subjects at both timepoints excluding voxels that were part of the original WMH mask so that no voxels were in both the lesionometric ROI and a WMH (for individual examples see Fig. 2).

**Figure 2:**
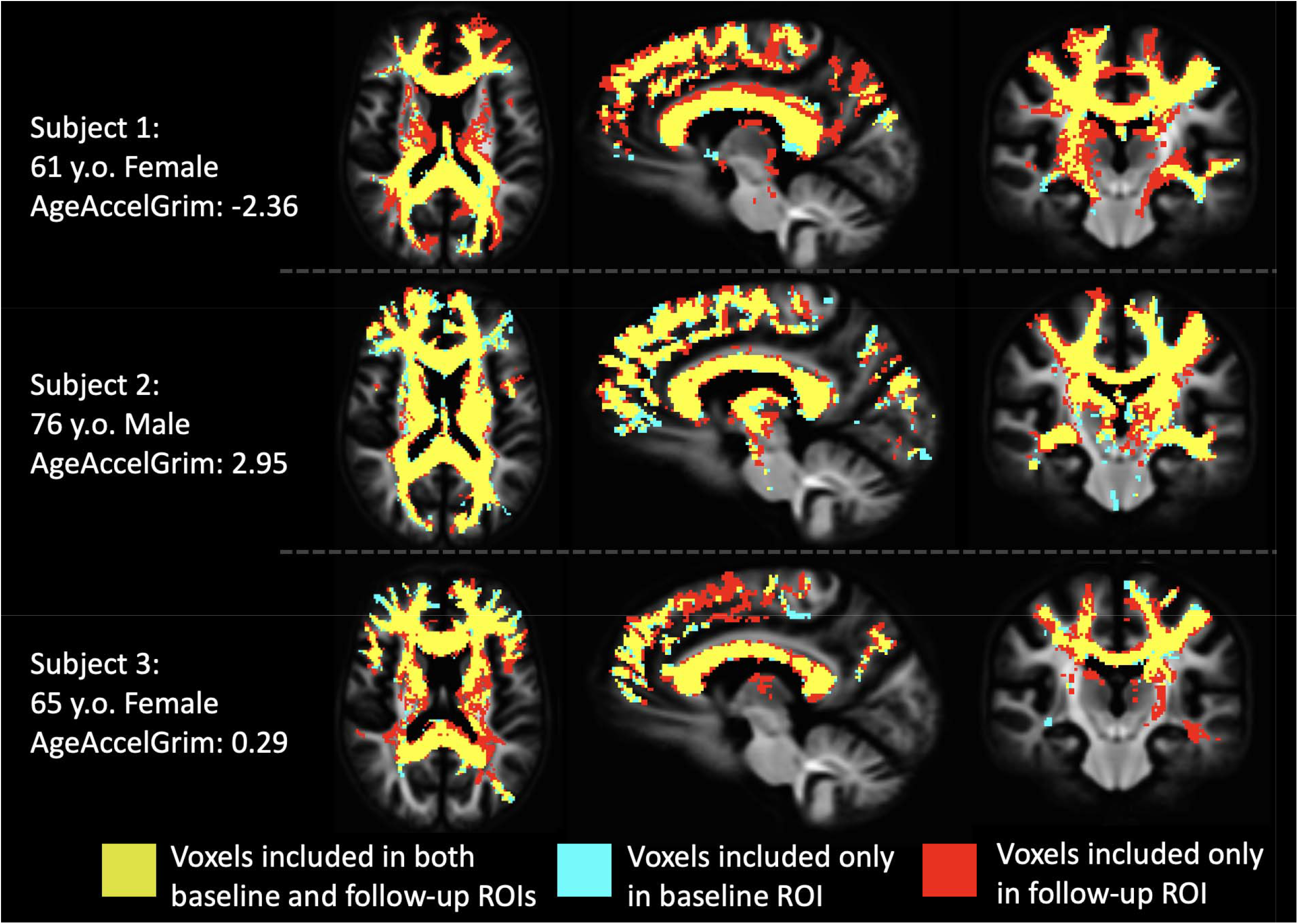
Illustration of subject-specific lesionometry ROIs in the whole-group template space from 3 example participants. Each subject in the study at each timepoint contributed a unique scan- and subject-specific ROI for analysis. ROIs were generated by filtering any ‘lesioned’ axonal tracts from the whole brain tractogram that passed through a voxel identified as being part of a WMH. This WMH-derived tractogram was then converted to a typical binary voxel ROI with the use of a low pass filter to only include voxels containing 10 or more tracts to ensure consistency. The volume of the ROI once the WMH volume was corrected for was not significantly different between baseline and follow-up (F_1,36_=0.0272, p=0.870 n.s.).

### Epigenetic Analysis

For full description of the epigenetic protocol see supplementary methods. Briefly, 8.5 ml blood were drawn from each participant at the baseline visit and DNA was extracted and amplified with PCR before being assayed using the Illumina Infinium MethylationEPIC BeadChip according to manufacturer instruction. The R packages *minfi* and *shinyMethyl* were used for background subtraction, dye-bias normalization, removal of missing values, quality control, and to check for batch effects^45–49^. All samples passed Illumina quality controls as assessed using the *ewastools* R package^50^. Unnormalized betas were filtered to include CpGs specified by Horvath as necessary for calculation of various clocks^1^. The betas were uploaded to Horvath’s online DNA methylation age calculator (htpps://dnamage.genetics.ucla.edu), which provides measures of DNA methylation GrimAge^3^. AgeAccelGrim was calculated by regression of GrimAge onto subject age, providing a chronological age normed value of accelerated or decelerated mortality risk relative to the subject’s age at baseline.

### Statistical Analysis

After AgeAccelGrim was calculated for each subject at baseline a general linear model was constructed using a planned set of covariates to test their association with GrimAge. Subject chronological age at baseline, sex, and total brain volume at baseline were initially tested for a relationship with AgeAccelGrim. Missing longitudinal data was analyzed to determine whether data are missing completely at random. Missingness was not associated with chronological age at scan acquisition (T_1,96_=-1.141, p=0.257 n.s.), sex (T_1,96_=1.535, p=0.128 n.s.), total brain volume (T_1,96_=-0.886, p=0.3778 n.s.), WMH volume (T_1,96_=-0.258, p=0.797 n.s.), GrimAge (T_1,96_=-0.706, p=0.482 n.s.) or AgeAccelGrim (T_1,96_=0.349, p=0.728 n.s.). Following this analysis for all imaging results unless otherwise noted, chronological age at scan acquisition, sex, and a volumetric component of either the whole brain or the subject- and scan-specific ROI were used as covariates.

For longitudinal results, a within-subjects ANOVA approach was used to specifically examine the 41 subjects scanned longitudinally, with controls for subject sex, age at baseline, and total brain volume at both timepoints when appropriate.

## Results

### AgeAccelGrim is associated with sex and brain volume

As expected, there was no significant relationship between AgeAccelGrim and chronological age in the baseline sample (T_4,94_=-0.431, p=0.667 n.s.). A highly significant relationship between AgeAccelGrim and sex (T_4,94_=-5.200, p<0.001) was also expected, given that sex is included in GrimAge calculation^3^. Interestingly, female subjects typically have a lower AgeAccelGrim, particularly after 65 years of age. There was also a negative relationship between AgeAccelGrim and total brain volume (T_4,94_=-2.782, p<0.01) when controlled for age and sex.

### AgeAccelGrim is not associated with whole brain dMRI microstructure metrics

Significance values that follow are for AgeAccelGrim as a predictor of the respective microstructural signal fraction. Whole brain microstructural composition was first assessed at baseline for relationships to AgeAccelGrim. There was no significant relationship between global ICI signal fraction (T_5,93_=1.353, p=0.179 n.s.), nor global ICA signal fraction (T_5,93_=-0.298, p=0.766 n.s.), but there was a trend toward significance with global ECI signal fraction (T_5,93_=1.797, p=0.0755 n.s.) indicating that AgeAccelGrim may be signaling the presence of extracellular water in the aging brain (Fig. 3a).

**Figure 3:**
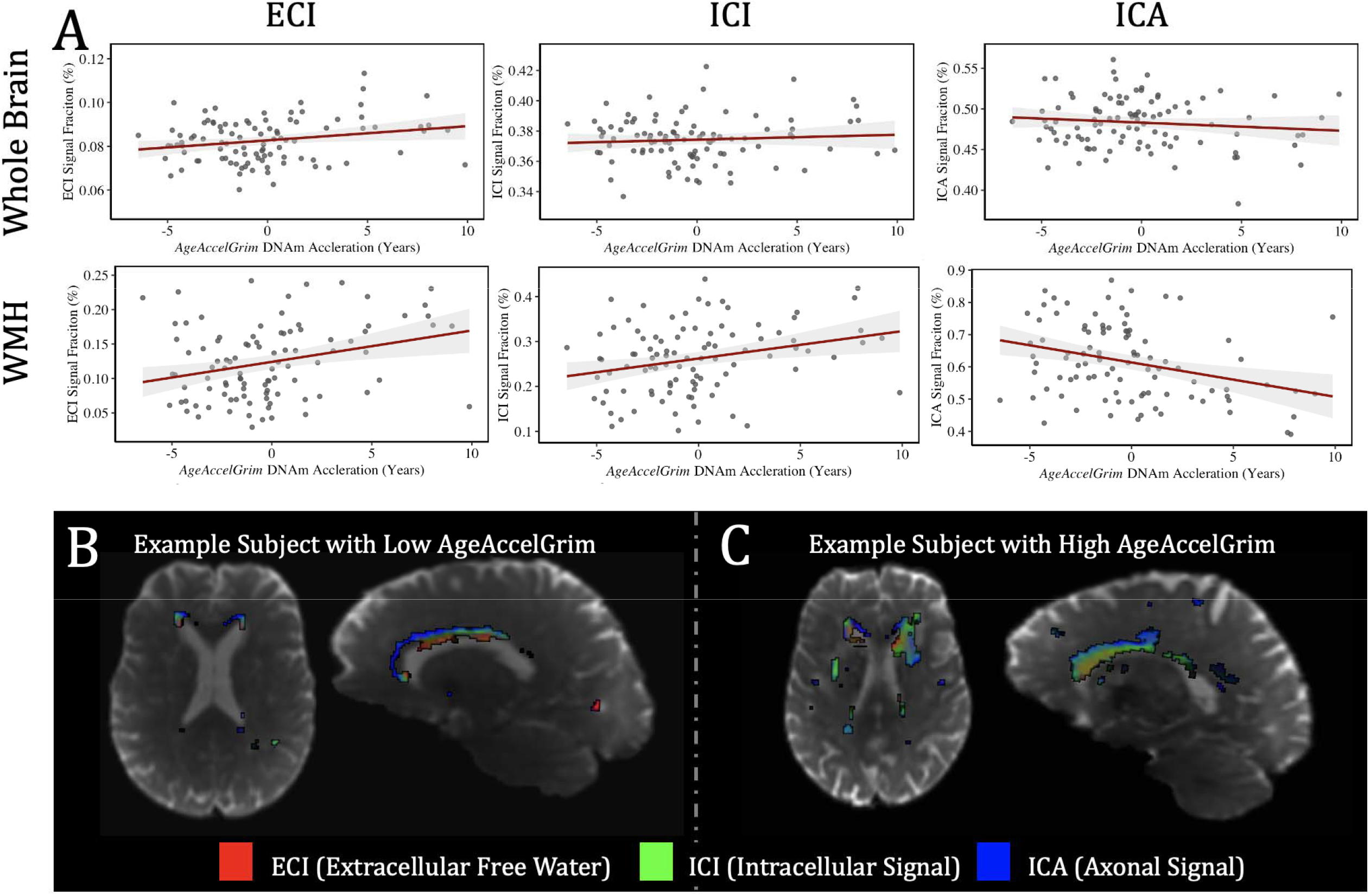
(A) Charts showing the relationship between whole brain (top row) and WMH (bottom row) 3T-CSD microstructure measurements (from left to right: ECI, ICI, and ICA signal fractions) and AgeAccelGrim at baseline. All three microstructural tissue compartments averaged across the whole brain did not have a significant relationship with AgeAccelGrim ECI: T_6,93_=1.797, p=0.076 n.s.; ICI: T_6,93_=1.353, p=0.179 n.s.; ICA: T_6,93_=-0.298, p=.767 n.s.). But when measured exclusively within the WMH all microstructure compartments had a significant relationship with AgeAccelGrim (ECI: T_6,92_=2.844, p<0.01; ICI: T_6,92_=2.741, p<0.01; ICA: T_6,92_=-3.140, p<0.01). (B) Image of an example subject with low AgeAccelGrim with voxels composing the WMH RGB color-coded based on the respective proportion of signal fraction composition (ECI in red, ICI in green, and ICA in blue). (C) Image of an example subject with high AgeAccelGrim with voxels composing the WMH colored using the same approach. The subject with high AgeAccelGrim shows characteristically elevated levels of ECI and ICI signal fraction throughout the area identified as belonging to a WMH while in the low AgeAccelGrim subject the WMH is still largely composed of ICA signal fraction, indicating that its composition is still similar to healthy nearby WM.

### AgeAccelGrim predicts WMH size and microstructural composition at baseline

Moving into subject-specific analysis of WMH, 3T-CSD microstructure and volume were measured within each subjects’ WMH at baseline. Greater WMH volume was significantly predicted by increased AgeAccelGrim (T_5,93_=2.931, p<0.01) and was added as an additional volumetric control variable for the WMH microstructure model to account for variation in WMH volume between subjects. All three microstructural tissue compartments averaged across the WMH had a significant relationship with AgeAccelGrim (ECI: T_6,92_=2.844, p<0.01; ICI: T_6,92_=2.741, p<0.01; ICA: T_6,92_=-3.140, p<0.01). For these models ECI and ICA had additional significant relationships with WMH volume (ECI: T_6,92_=5.537, p<0.001; ICA: T_6,92_=-2.494, p<0.05) while ICI did not have a significant relationship with WMH volume (ICI: T_6,92_=0.348, p=0.728 n.s.) suggesting that spatial location of the WMH plays a role in tissue composition but that AgeAccelGrim is able to predict ICI signal fraction and increased ECI signal fraction, occurring at the expense of ICA signal fraction (Fig. 3b).

### AgeAccelGrim does not predict WMH size and microstructural composition longitudinally

Looking longitudinally however AgeAccelGrim collected at baseline was less predictive of rates of changes in these metrics. As expected in an aging cohort, total brain volume significantly declined between baseline and follow-up scans (F_1,40_=9.163, p<0.01), while WMH volume significantly increased (F_1,40_=7.688, p<0.01). For the microstructural metrics between baseline and follow-up whole brain ECI signal fraction significantly increased (F_1,36_=5.395, p<0.05), while significant decreases were observed in whole brain ICI signal fraction (F_1,36_=4.075, p<0.05) and no significant change was observed in ICA signal fraction (F_1,36_=1.776, p=0.189 n.s.). AgeAccelGrim was not significantly predictive of changes in total brain volume (F_1,37_=3.172, p=0.083 n.s.) (Fig. 4) nor was it predictive of any changes in whole brain cellular microstructure measurements (ECI: F_1,36_= 0.412, p=0.525 n.s.; ICI: F_1,36_=0.173, p=0.680 n.s.; ICA: F_1,36_=1.292, p=0.263 n.s.). Despite strong correlations in the baseline data AgeAccelGrim was also not predictive of changes in WMH volume (F_1,36_=2.377, p=0.131 n.s.) nor any microstructural composition measures (ECI: F_1,36_=1.738, p=0.195 n.s.; ICI: F_1,36_=0.141, p=0.709 n.s.; ICA: F_1,36_=0.861, p=0.359 n.s.) (Fig. 5).

**Figure 4:**
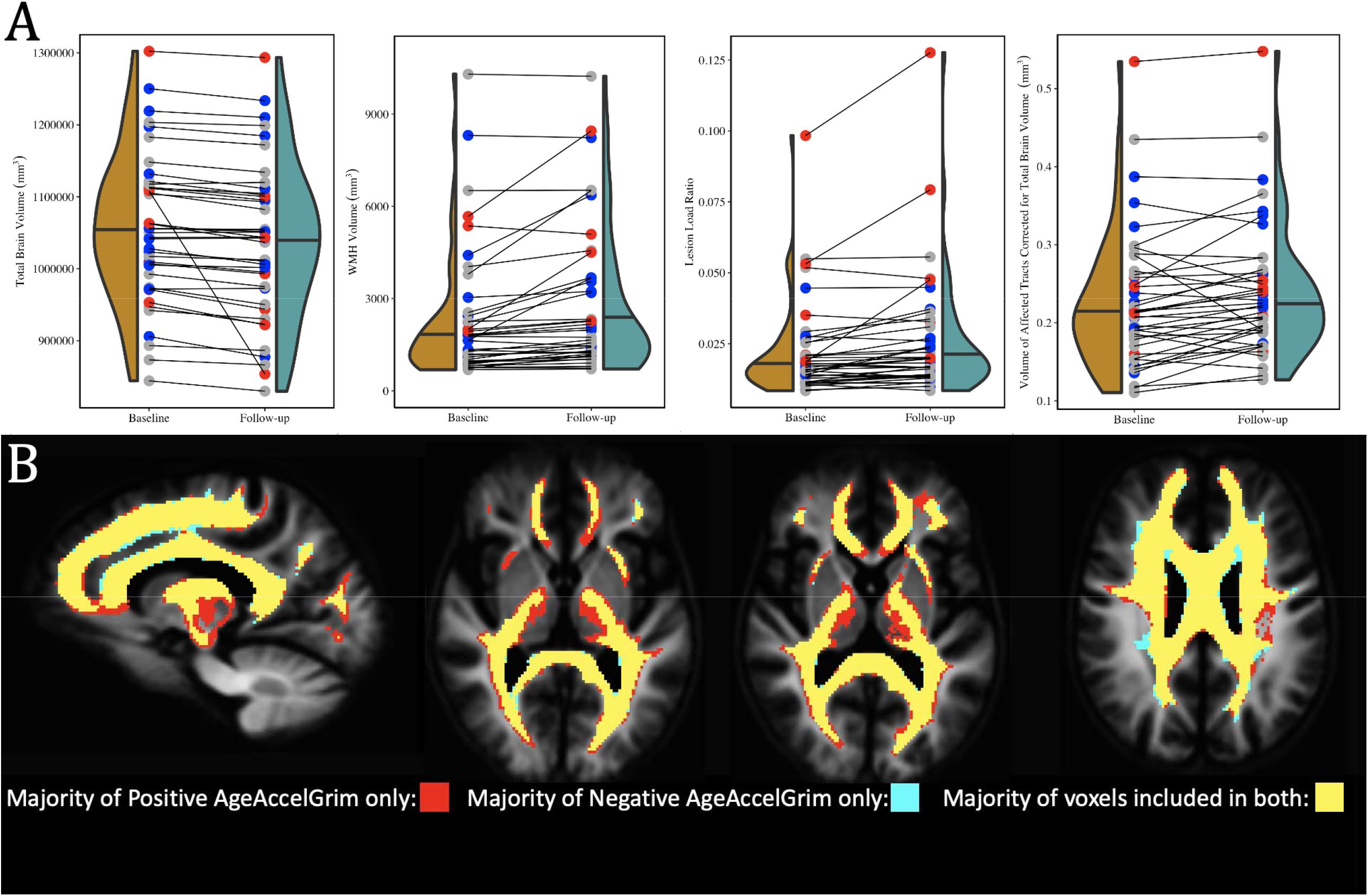
(A) Charts displaying the observed longitudinal change in total brain volume, WMH volume, lesion load ratio (WMH volume to lesionometry ROI volume), and the volume of the lesionometry ROI volume corrected for total brain volume, from left to right. Individual subjects are represented in each chart by points at both baseline and follow-up, and are colored according to AgeAccelGrim measured at baseline (Blue = low AgeAccelGrim (less than -2), Red = high AgeAccelGrim (greater than 2), Gray = AgeAccelGrim close to chronological age (between -2 and 2)). Longitudinally, AgeAccelGrim was not significantly predictive of changes in total brain volume (F_1,37_=3.172, p=0.083 n.s.) and was also not predictive of changes in WMH volume (F_1,36_=2.377, p=0.131 n.s.). AgeAccelGrim was also did not significantly predict the longitudinal change in size of the network passing through the WMH (F_1,36_=3.476, p=0.070 n.s.) but had a significant relationship with lesion load (F_1,36_=5.397, p<0.05). (B) Images displaying the overlapping locations included in the lesionometry ROIs. The baseline lesionometry ROIs in template space from each subject were divided into two groups depending on AgeAccelGrim, with the positive group having a value greater than 0 indicating accelerated aging and the negative group having a value lower than 0 indicating slowed aging. A voxel was included in that group’s mask if it was present in a majority (>50%) of subject’s lesionometry ROIs. Both groups masks are presented above with voxels unique to each colored respectively (red for positive and blue for negative) and voxels common to both colored in yellow. Positive AgeAccelGrim subjects were more likely to have affected tracts that extended into the thalamus and frontal lobe, while negative AgeAccelGrim subjects were more likely to have affected periventricular and cingulate tracts.

**Figure 5:**
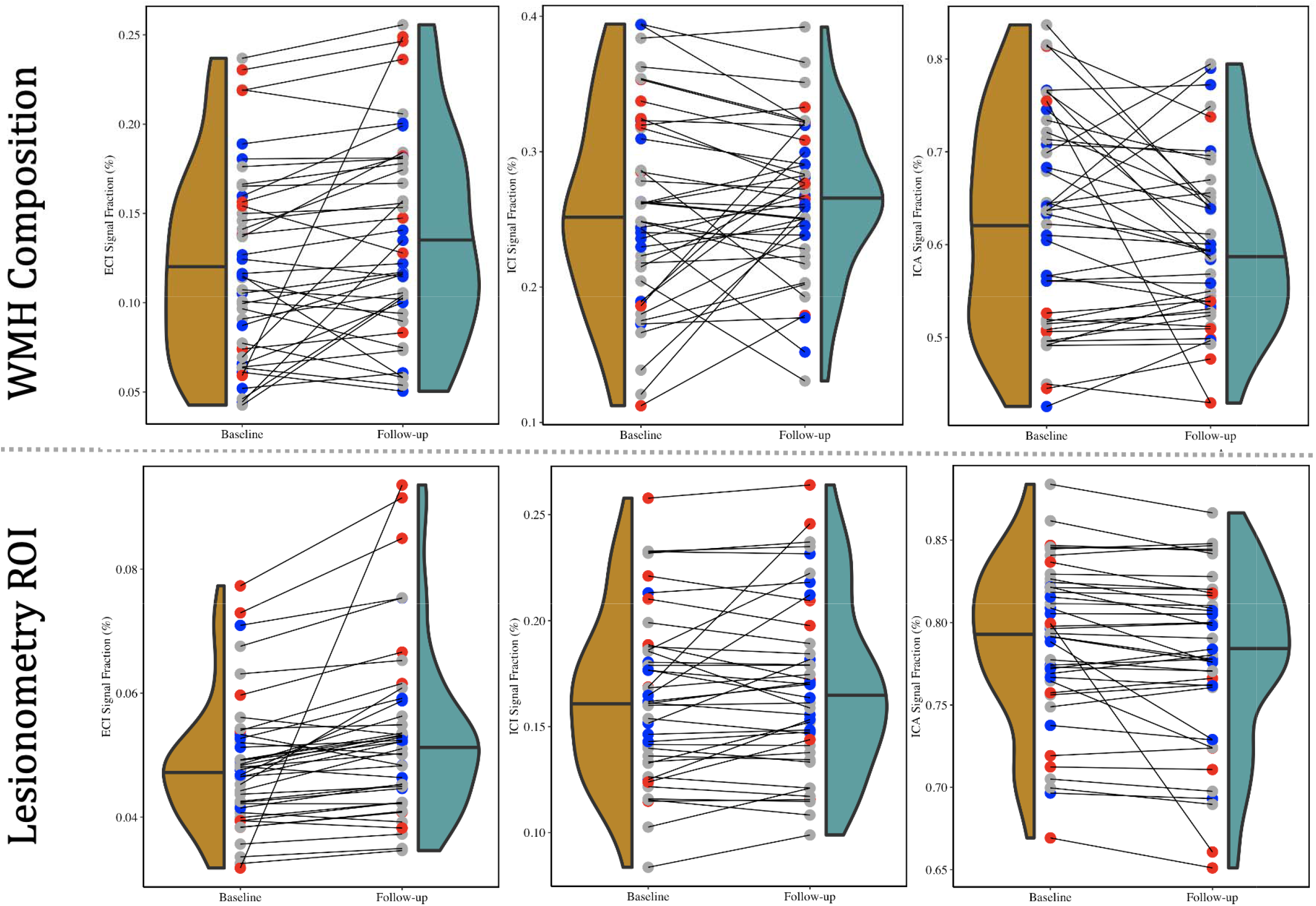
Charts displaying longitudinal 3T-CSD microstructural results from both WMH and lesionometry ROIs from each of the 3 signal fraction compartments (ECI, ICI, and ICA, arranged left to right). Individual subjects are colored according to AgeAccelGrim measured at baseline (Blue = low AgeAccelGrim (less than -2), Red = high AgeAccelGrim (greater than 2), Gray = AgeAccelGrim close to chronological age (between -2 and 2)). Despite strong correlations in the baseline data AgeAccelGrim was not predictive of longitudinal changes in any WMH microstructural composition measures (ECI: F_1,36_=1.738, p=0.195 n.s.; ICI: F_1,36_=0.141, p=0.709 n.s.; ICA: F_1,36_=0.861, p=0.359 n.s.). However AgeAccelGrim in the lesionometry ROI was able to significantly predict longitudinal change in each signal fraction compartment, with a positive relationship between AgeAccelGrim and ECI signal fraction (F_1,36_=11.11, p<0.01), a positive relationship between AgeAccelGrim and ICI signal fraction (F_1,36_=4.353, p<0.05), and a negative relationship between AgeAccelGrim and ICA signal fraction (F_1,36_=6.243, p<0.05).

### AgeAccelGrim predicts baseline and longitudinal changes in microstructural composition in lesionometry ROIs

The microstructure measurements taken from the lesionometry ROIs, were able to be predicted by AgeAccelGrim both cross-sectionally at baseline and longitudinally. This contrasts with the lack of longitudinal relationships between AgeAccelGrim and microstructure both at whole brain and within the WMH. At baseline AgeAccelGrim was trending toward a significant positive relationship with the size of the network passing through the WMH (volume of lesionometry ROI; T_5,93_=1.976, p=0.0512) and was not significantly predictive of longitudinal change (F_1,36_=3.476, p=0.070 n.s.) (Fig. 4). The subsequent microstructural models are corrected for subject age at baseline, sex, and volume of the lesionometry ROI. At baseline AgeAccelGrim had a significantly positive relationship with ECI signal fraction in the lesionometric ROI (T_5,93_=2.586, p<0.05), a positive relationship with ICI signal fraction (T_5,93_=2.073, p<0.05), and a negative relationship with ICA signal fraction (T_5,93_=-2.299, p<0.05). Longitudinally AgeAccelGrim was able to significantly predict the change in microstructural measurements in the lesionometry ROI between baseline and follow-up for each signal fraction compartment (Fig. 5), with a positive relationship between AgeAccelGrim and ECI signal fraction (F_1,36_=11.11, p<0.01), a positive relationship between AgeAccelGrim and ICI signal fraction (F_1,36_=4.352, p<0.05), and a negative relationship between AgeAccelGrim and ICA signal fraction (F_1,36_=6.243, p<0.05). This is particularly interesting because when AgeAccelGrim is removed and the lesionometry ROIs are exclusively tested for longitudinal change between scanning sessions (including sex, age at baseline, and ROI volume as controls identical to before) there was only a significant difference between baseline and follow-up for the ECI signal fraction (F_1,37_=8.846, p<0.01) and there was no significant difference between baseline and follow-up for the ICI signal fraction (F_1,37_=1.143, p=0.291 n.s.) nor for the ICA signal fraction (F_1,37_=4.051, p=0.051 n.s.). Finally, AgeAccelGrim had a positive relationship with lesion load, a ratio between the volume of each subject’s WMH and the volume of the lesionometry ROI (which does not include the WMH) at baseline (T_5,93_=4.245, p<0.001) and longitudinally (F_1,36_=5.397, p<0.05) (Fig. 4).

## Discussion

By examining markers of SVD using advanced measures of diffusion microstructure in a subject-specific lesionometry approach, this study has established a connection between a blood-based measure of mortality risk and neuronal damage. AgeAccelGrim was significantly correlated with WMH volume, supporting previous findings from other groups^15^ and was significantly correlated with WMH microstructural composition. To date, this study is the first to demonstrate an association between accelerated epigenetic age derived from peripheral blood and brain cellular microstructure^16–18^. Specifically, AgeAccelGrim was associated with higher WMH ECI or extracellular free water signal fraction, higher ICI signal fraction, and lower ICA or axonal signal fraction (which in healthy WM areas can be typically observed at values exceeding 90-95% signal fraction). This either indicates that cellular microstructure in WMH areas was more heavily damaged in individuals with accelerated epigenetic age, that that WMH in these subjects was located in a specific spatial periventricular arrangement^23^, or a combination of both (Fig. 3b&c, Fig. 4b).

Analyzing the axonal bundles that passed through the WMH showed that AgeAccelGrim was able to predict the size of the lesionometric ROI, an increased lesion burden, and increased degeneration within the lesionometric ROI. The subject-specific lesionometric approach combined with 3T-CSD analysis of cellular microstructure was able to isolate areas of the brain vulnerable to SVD-related damage. These findings are further reinforced by the lack of longitudinal association between accelerated DNAm and whole brain microstructural measurements. It is necessary to narrowly define localized regions of the brain where vascular damage is likely occurring at an individual level. Related to this idea is the observed significant relationship between AgeAccelGrim and increased ICI signal fraction. When 3T-CSD is applied to the brain the ICI signal fraction (also referred to as the GM-like signal fraction) predominates in the cortex. Observing increased ICI signal fraction, especially within the WM skeleton where the WMH and lesionometry ROIs are located, is possibly indicative of increased neuroinflammation or activated glial cells in response to injury^51^. The increased ECI signal fraction however, is straightforwardly interpretable as either edema or the absence of cellular tissue as a result of axonal degeneration. Together these longitudinal microstructure results indicate that AgeAccelGrim can predict subsequent neuronal deterioration over multiple years. This suggests that GrimAge may be a useful marker for positioning an individual on the trajectory of age-related neuronal decline, mediated via cardiovascular factors and SVD.

This study did not address the potential change in GrimAge calculation between baseline and follow-up, though the clock is better suited for predictive use and it not well established how GrimAge varies over time. Due to screening procedures within the VCAP cohort examined here, it is unlikely that participants would have undertaken a major lifestyle change that would have altered their AgeAccelGrim, such as beginning a prolific smoking habit, between the baseline and follow-up recruitment. Additionally, the relatively short period of time would likely not be long enough to impart significant change, given that a study using multiple clocks, including Horvath’s and Hannum’s, showed that age-related increases in epigenetic age tend to flatten out in older age along a logarithmic trend^52^. Follow-up work to this study will aim to tie performance on cognitive tasks performed during assessment to epigenetic, cardiovascular, and diffusion microstructure metrics to evaluate the behavioral output of changed observed here. Further refinement of the GrimAge clock could include different combinations of plasma-proteins in order to discern which proteins are primarily driving observed changed in brain microstructure instead of general mortality.

The degree to which these results are based in WMH suggests a cardiovascular connection via SVD between the epigenetic clock estimates provided by AgeAccelGrim and brain cellular microstructure. While GrimAge has generally been shown to be highly predictive of the development of several cardiovascular health related pathologies such as time-to-coronary heart disease, congestive heart failure, hypertension, type 2 diabetes, and physical functioning^3^, it is still unknown exactly which features of the cardiovascular system drive this change and contribute to the results seen in this study. Several other studies have found that GrimAge is related to heart failure^53^ and composite measures of whole cardiovascular health including diet, smoking, physical activity body mass index, blood pressure, total cholesterol, and blood glucose^54^ but physiological measures of brain cardiovascular health have not been clearly associated with GrimAge^55^. WMH volume has previously been used as a biomarker for SVD severity^7^ and ECI signal fraction analogues such as free water have been established as a marker for cerebral SVD^56^. These studies suggest that the results presented in this study indicate that AgeAccelGrim may be a biomarker for SVD-related brain injury and degeneration.

This study has provided evidence that a blood-based epigenetic marker of age acceleration can predict the degenerative effects of SVD in the brain. AgeAccelGrim was able to predict the volume and composition of WMH as well as widespread diffusion microstructure signatures of neuronal decline in a subject-specific manner.

## Abbreviations

MRI: Magnetic Resonance Imaging
WMH: White Matter Hyperintensity
SVD: Small Vessel Disease
3T-CSD: 3-Tissue Constrained Spherical Deconvolution
DNAm: DNA methylation
VCAP: Virginia Cognitive Aging Project
WM: White Matter
GM: Gray Matter
CSF: Cerebrospinal Fluid
FOD: Fiber Orientation Distribution
SS3T-CSD: Single-Shell 3-Tissue Constrained Spherical Deconvolution
ROI: Region of Interest
ECI: Extracellular Isotropic
ICI: Intracellular Isotropic
ICA: Intracellular Anisotropic

## Supplementary Methods

### Epigenetic Age

Eight and a half milliliters of whole blood were drawn into a PAXgene Blood DNA Tube (PreAnalytiX, Hombrechtikon, Switzerland). Samples were stored at 20°C for short-term storage (up to 3 months) then transferred to -80°C for long-term storage. DNA was extracted using the PAXgene Blood DNA kit (PreAnalytiX, Hombrechtikon, Switzerland) according to manufacturer instructions. DNA concentration was determined by Quant-iT™ PicoGreen® dsDNA reagent (Thermofisher Scientific, Waltham, MA, USA) per manufacturers instruction. Florescence was detected using a Tecan Infinite M200 Pro microplate reader (Tecan, Switzerland). 500 ng of DNA was bisulfite treated using a Zymo EZ DNA Methylation kit (Zymo Research, Irvine, CA) using the following PCR conditions for Illumina’s Infinium Methylation assay (95°C for 30 seconds, 50°C for 60 minutes×16 cycles). DNA methylation was assayed using the Illumina Infinium MethylationEPIC BeadChips. A total of 4μL of bisulfite converted DNA was hybridized to Illumina BeadChips according to the manufacturer’s protocols. Samples were denatured and amplified overnight for 20 to 24 hours. After overnight incubation, fragmentation, precipitation, and resuspension of the samples. Samples were then hybridized to EPIC BeadChips for 16 to 24 hours. BeadChips were washed to remove any unhybridized DNA and labeled with nucleotides to extend the primers to the DNA sample. BeadChips were imaged using the Illumina iScan system (Illumina) according to the Infinium HD methylation protocol.

Processing of methylation array data was completed as previously reported^57^. Raw .idat files were read and preprocessed using the minfi R package^45,47^. The data set was preprocessed using noob for background subtraction and dye-bias normalization. All methylation values with detection P>0.01 were set to missing (median sample: 765 probes, range: 319 to 4453), and probes with >1% missing values (n=6,663) were removed from further analysis. All samples were checked and confirmed to ensure that predicted sex matched reported sex. Additionally, samples were checked for excessive missing data (>5%) and unusual cell mixture estimates, which was estimated using the Houseman method as implemented in minfi^48,49^. All samples passed these quality controls. Principal components analysis, as implemented in the shinyMethyl R package, was used to examine batch effects ^46^. The first seven principal components were examined using plots and potential batch effects were tested using linear models. Principal components 3 and 6, which account for 2.38% and 1.65% of total variance respectively, were associated with position on the array (PC3: F_(7, 100)_ = 6.668, p = 1.77e-6, adjusted R^2^ = 0.271; PC6: F_(7, 100)_ = 2.328, p = 0.030, adjusted R^2^ = 0.080). Principal components 1, 4, and 5, which account for 3.63%, 1.89%, and 1.77% of the total variance were associated with bisulfite conversion plate (PC1: F_(1, 106)_ = 9.918, p = 0.002, adjusted R^2^ = 0.077; PC4: F_(1, 100)_ = 34.04, p = 5.932e-8, adjusted R^2^ = 0.236; PC5: F_(1, 100)_ = 31.07, p = 1.91e-7, adjusted R^2^ = 0.219). Principal components 4 and 5, were associated with array (PC4: F_(13, 94)_ = 4.332, p = 1.14e-5, adjusted R^2^ = 0.288; PC5: F_(13, 94)_ = 4.229, p = 1.06e-5, adjusted R^2^ = 0.282). Bisulfite conversion plate and array number were associated with each other, as samples on the same array originated from the same bisulfite conversion plate. Because samples were randomized across plates and arrays, and proportions of variance explained by associated principle components were low, no batch correction method was used. The ewastools R package was used to assess Illumina quality control metrics and call genotypes and donor IDs to ensure the identity of repeated samples from the same individual^50^. All samples passed Illumina quality controls.

To determine assay variability, we included one set of five technical replicates and an additional three sets of two technical replicates. After quality control filters and normalization procedures were applied, the 5,000 CpGs with the most variable M values were used as input for calculating Pearson’s correlation coefficients among all pairwise combinations of samples.

Pearson’s correlation of unrelated samples (different individuals) were below 0.8. Pearson’s while correlations of technical replicates ranged from 0.988-0.994, indicating high agreement between technical replicates. Unnormalized betas were filtered to include CpGs specified by Horvath as necessary for calculation of various clocks. The betas were uploaded to Horvath’s online DNA methylation age calculator (htpps://dnamage.genetics.ucla.edu), which provides measures of Horvath’s multi-tissue age estimator^1^, DNA methylation GrimAge^3^, and cell type abundance. A sample annotation file was included. The options to normalize data and apply advanced analysis were selected. Technical replicates were used to determine measurement error of DNAmAge, the output of Horvath’s multi-tissue age estimator. The absolute difference of DNAmAge between technical replicate pairs was taken, as was the highest absolute difference in the set of five technical replicates. The median of the absolute difference was 2.02 years (range: 0.44-5.73 years).

## Notes

### Competing Interest Statement

The authors have declared no competing interest.

## Works Cited

1. Horvath S. DNA methylation age of human tissues and cell types. Genome biology. 2013;14(10):1–20.

2. Lin Q, Weidner CI, Costa IG, et al. DNA methylation levels at individual age-associated CpG sites can be indicative for life expectancy. Aging (Albany NY). 2016;8(2):394.

3. Lu AT, Quach A, Wilson JG, et al. DNA methylation GrimAge strongly predicts lifespan and healthspan. Aging (Albany NY). 2019;11(2):303.

4. Felder RB, Francis J, Zhang ZH, Wei SG, Weiss RM, Johnson AK. Heart failure and the brain: new perspectives. American Journal of Physiology-Regulatory, Integrative and Comparative Physiology. 2003;284(2):R259–R276.

5. Hillman CH, Erickson KI, Kramer AF. Be smart, exercise your heart: exercise effects on brain and cognition. Nature reviews neuroscience. 2008;9(1):58–65.

6. Jefferson AL, Himali JJ, Beiser AS, et al. Cardiac index is associated with brain aging: the Framingham Heart Study. Circulation. 2010;122(7):690–697.

7. Finsterwalder S, Vlegels N, Gesierich B, et al. Small vessel disease more than Alzheimer’s disease determines diffusion MRI alterations in memory clinic patients. Alzheimer’s & Dementia. 2020;16(11):1504–1514.

8. Tuladhar AM, van Norden AG, de Laat KF, et al. White matter integrity in small vessel disease is related to cognition. NeuroImage: Clinical. 2015;7:518–524.

9. Wardlaw JM, Smith C, Dichgans M. Small vessel disease: mechanisms and clinical implications. The Lancet Neurology. 2019;18(7):684–696.

10. Di Stadio A, Messineo D, Ralli M, et al. The impact of white matter hyperintensities on speech perception. Neurological Sciences. 2020;41(7):1891–1898.

11. Gunning-Dixon FM, Raz N. The cognitive correlates of white matter abnormalities in normal aging: a quantitative review. Neuropsychology. 2000;14(2):224.

12. Maillard P, Mitchell GF, Himali JJ, et al. Effects of arterial stiffness on brain integrity in young adults from the Framingham Heart Study. Stroke. 2016;47(4):1030–1036.

13. Duering M, Finsterwalder S, Baykara E, et al. Free water determines diffusion alterations and clinical status in cerebral small vessel disease. Alzheimer’s & Dementia. 2018;14(6):764–774.

14. Dawber TR, Meadors GF, Moore Jr FE. Epidemiological approaches to heart disease: the Framingham Study. American Journal of Public Health and the Nations Health. 1951;41(3):279–286.

15. Hillary RF, Stevenson AJ, Cox SR, et al. An epigenetic predictor of death captures multi-modal measures of brain health. Molecular psychiatry. Published online 2019:1–11.

16. Fransquet PD, Lacaze P, Saffery R, et al. Accelerated Epigenetic Aging in Peripheral Blood does not Predict Dementia Risk. Current Alzheimer Research. 2021;18(5):443–451.

17. Russ TC. DNA methylation-based measures of accelerated biological ageing and the risk of dementia in the oldest-old: a study of the Lothian Birth Cohort 1921. Published online 2020.

18. Shadyab AH, McEvoy LK, Horvath S, et al. Association of Epigenetic Age Acceleration With Incident Mild Cognitive Impairment and Dementia Among Older Women. The Journals of Gerontology: Series A. Published online 2021.

19. Biesbroek JM, Weaver NA, Hilal S, et al. Impact of strategically located white matter hyperintensities on cognition in memory clinic patients with small vessel disease. PLoS One. 2016;11(11):e0166261.

20. Chamberland M, Winter M, Brice TA, Jones DK, Tallantyre EC. Beyond lesion-load: tractometry-based metrics for characterizing white matter lesions within fibre pathways. Published online 2020.

21. Winter M, Tallantyre EC, Brice TA, Robertson NP, Jones DK, Chamberland M. Tract-specific MRI measures explain learning and recall differences in multiple sclerosis. Brain Communications. 2021;3(2):fcab065.

22. Khan W, Khlif MS, Mito R, Dhollander T, Brodtmann A. Investigating the microstructural properties of normal-appearing white matter (NAWM) preceding conversion to white matter hyperintensities (WMHs) in stroke survivors. NeuroImage. 2021;232:117839.

23. Mito R, Dhollander T, Xia Y, et al. In vivo microstructural heterogeneity of white matter lesions in healthy elderly and Alzheimer’s disease participants using tissue compositional analysis of diffusion MRI data. NeuroImage: Clinical. 2020;28:102479.

24. Salthouse TA. Trajectories of normal cognitive aging. Psychology and aging. 2019;34(1):17.

25. Siedlecki K. Findings From the Virginia Cognitive Aging Project: Individual Differences, Well-Being, and Cognition. Innovation in Aging. 2020;4(Supplement_1):591–592.

26. Gunter J, Thostenson K, Borowski B, et al. ADNI-3 MRI Protocol. Alzheimer’s Dementia. 2017;13(7):P104–P105.

27. Dhollander T, Connelly A. A novel iterative approach to reap the benefits of multi-tissue CSD from just single-shell (+ b= 0) diffusion MRI data. In: Proc ISMRM. Vol 24. ; 2016:3010.

28. Jeurissen B, Tournier JD, Dhollander T, Connelly A, Sijbers J. Multi-tissue constrained spherical deconvolution for improved analysis of multi-shell diffusion MRI data. NeuroImage. 2014;103:411–426. doi:10.1016/j.neuroimage.2014.07.061

29. Tournier JD, Smith R, Raffelt D, et al. MRtrix3: A fast, flexible and open software framework for medical image processing and visualisation. NeuroImage. 2019;202:116137.

30. Jenkinson M, Beckmann CF, Behrens TE, Woolrich MW, Smith SM. Fsl. Neuroimage. 2012;62(2):782–790.

31. Smith SM, Jenkinson M, Woolrich MW, et al. Advances in functional and structural MR image analysis and implementation as FSL. Neuroimage. 2004;23:S208–S219.

32. Veraart J, Fieremans E, Novikov DS. Diffusion MRI noise mapping using random matrix theory. Magnetic resonance in medicine. 2016;76(5):1582–1593.

33. Kellner E, Dhital B, Kiselev VG, Reisert M. Gibbs-ringing artifact removal based on local subvoxel-shifts. Magnetic resonance in medicine. 2016;76(5):1574–1581.

34. Andersson JL, Graham MS, Zsoldos E, Sotiropoulos SN. Incorporating outlier detection and replacement into a non-parametric framework for movement and distortion correction of diffusion MR images. Neuroimage. 2016;141:556–572.

35. Andersson JL, Sotiropoulos SN. An integrated approach to correction for off-resonance effects and subject movement in diffusion MR imaging. Neuroimage. 2016;125:1063–1078.

36. Fischl B. FreeSurfer. Neuroimage. 2012;62(2):774–781.

37. Avants BB, Tustison NJ, Stauffer M, Song G, Wu B, Gee JC. The Insight ToolKit image registration framework. Front Neuroinform. 2014;8. doi:10.3389/fninf.2014.00044

38. Dhollander T, Raffelt D, Connelly A. Unsupervised 3-tissue response function estimation from single-shell or multi-shell diffusion MR data without a co-registered T1 image. In: ISMRM Workshop on Breaking the Barriers of Diffusion MRI. Vol 5. ISMRM; 2016.

39. Newman BT, Untaroiu A, Druzgal TJ. A novel diffusion registration method with the NTU-DSI-122 template to transform free water signal fraction maps to stereotaxic space. 2020;Proceedings of the ISMRM 28th General Meeting.

40. Tournier JD, Calamante F, Connelly A. Improved probabilistic streamlines tractography by 2nd order integration over fibre orientation distributions. In: Proceedings of the International Society for Magnetic Resonance in Medicine. Vol 1670. John Wiley & Sons, Inc. New Jersey, USA; 2010.

41. Smith RE, Tournier JD, Calamante F, Connelly A. SIFT: Spherical-deconvolution informed filtering of tractograms. Neuroimage. 2013;67:298–312.

42. Newman BT, Dhollander T, Reynier KA, Panzer MB, Druzgal TJ. Test–retest reliability and long-term stability of three-tissue constrained spherical deconvolution methods for analyzing diffusion MRI data. Magn Reson Med. 2020;84(4):2161–2173. doi:10.1002/mrm.28242

43. Blair J C, Newman BT, Druzgal TJ. Evaluating Lifespan Tissue Structure: Comparing CSD Signal Fraction and VBM Grey Matter Density. Proceedings of the ISMRM 27th General Meeting. Published online 2019.

44. Hotz I, Deschwanden PF, Liem F, et al. Performance of Three Freely Available Methods for Extracting White Matter Hyperintensities: FreeSurfer, UBO Detector, and BIANCA. Wiley Online Library; 2022.

45. Aryee MJ, Jaffe AE, Corrada-Bravo H, et al. Minfi: a flexible and comprehensive Bioconductor package for the analysis of Infinium DNA methylation microarrays. Bioinformatics. 2014;30(10):1363–1369.

46. Fortin JP, Fertig E, Hansen K. shinyMethyl: interactive quality control of Illumina 450k DNA methylation arrays in R. F1000Research. 2014;3.

47. Fortin JP, Triche Jr TJ, Hansen KD. Preprocessing, normalization and integration of the Illumina HumanMethylationEPIC array with minfi. Bioinformatics. 2017;33(4):558–560.

48. Houseman EA, Accomando WP, Koestler DC, et al. DNA methylation arrays as surrogate measures of cell mixture distribution. BMC bioinformatics. 2012;13(1):1–16.

49. Jaffe AE, Irizarry RA. Accounting for cellular heterogeneity is critical in epigenome-wide association studies. Genome biology. 2014;15(2):1–9.

50. Heiss JA, Just AC. Identifying mislabeled and contaminated DNA methylation microarray data: an extended quality control toolset with examples from GEO. Clinical epigenetics. 2018;10(1):1–9.

51. Newman BT, Untaroiu A, Druzgal TJ. A method for post-surgical evaluation of targeting accuracy in transcranial focused ultrasound thalamic ablation. 28th International Society of Magnetic Resonance in Medicine. 2020;28:4111.

52. Snir S, Farrell C, Pellegrini M. Human epigenetic ageing is logarithmic with time across the entire lifespan. Epigenetics. 2019;14(9):912–926.

53. Nguyen S, Northuis CA, Guan W, et al. Epigenetic Clocks And Incident Heart Failure: The Atherosclerosis Risk In Communities (aric). Circulation. 2021;143(Suppl_1):A033–A033.

54. Joyce BT, Gao T, Zheng Y, et al. Epigenetic Age Acceleration Reflects Long-Term Cardiovascular Health. Circulation research. 2021;129(8):770–781.

55. Grodstein F, Lemos B, Yu L, et al. The association of epigenetic clocks in brain tissue with brain pathologies and common aging phenotypes. Neurobiology of Disease. Published online 2021:105428.

56. Huang P, Zhang R, Jiaerken Y, et al. White Matter Free Water is a Composite Marker of Cerebral Small Vessel Degeneration. Translational Stroke Research. Published online 2021:1–9.

57. Abdulrahim JW, Kwee LC, Grass E, et al. Epigenome-Wide Association Study for All-Cause Mortality in a Cardiovascular Cohort Identifies Differential Methylation in Castor Zinc Finger 1 (CASZ 1). Journal of the American Heart Association. 2019;8(21):e013228.

